# Layer-specific mitochondrial diversity across hippocampal CA2 dendrites

**DOI:** 10.1101/2021.12.28.474366

**Authors:** Katy Pannoni, Daniela Gil, Mikel Cawley, Logan Campbell, Shannon Farris

## Abstract

CA2 is an understudied subregion of the hippocampus that is critical for social memory. Previous studies identified multiple components of the mitochondrial calcium uniporter (MCU) complex as selectively enriched in CA2, however the functional significance of this enrichment remains unclear. The MCU complex regulates calcium entry into mitochondria, which in turn regulates mitochondrial transport and localization to active synapses. We found that MCU is strikingly enriched in CA2 distal apical dendrites, precisely where CA2 neurons receive entorhinal cortical input carrying social information. Further, MCU-enriched mitochondria in CA2 distal dendrites are larger compared to mitochondria in CA2 proximal apical dendrites and neighboring CA1 apical dendrites, an effect also seen with genetically labeled mitochondria and electron microscopy. MCU overexpression in neighboring CA1 led to larger mitochondria preferentially in proximal dendrites compared to distal dendrites and controls, suggesting that MCU may act as a coincidence detector linking synaptic activity to mitochondrial morphology and function. Our findings demonstrate that mitochondria are molecularly and structurally diverse across hippocampal cell types and circuits, and implicate MCU expression in regulating mitochondrial mass and layer-specific dendritic localization. Functionally distinct mitochondria in CA2 distal dendrites may confer unique synaptic and circuit properties underlying CA2 function in social memory.

## INTRODUCTION

Mitochondria are critical organelles that are responsible for a variety of functions in the cell, including generating energy, regulating calcium homeostasis, and signaling apoptosis (Duchen 2000; Vakifahmetoglu-Norberg, Ouchida, and Norberg 2017). There is mounting evidence of mitochondrial diversity across tissues and cell types and the functional significance of this heterogeneity is an area of active investigation (Pekkernaz and Wang 2022; Sprenger and Langer 2019; Fecher et al. 2019). In neurons, mitochondria are important for synaptic transmission, maintenance, and plasticity (Devine and Kittler 2018; Tang and Zucker 1997; Li et al. 2004; Stowers et al. 2002). Mitochondria are commonly found in axon terminals, as well as other sites with high energetic demand, and the loss of presynaptic mitochondria results in defective synaptic transmission (Stowers et al. 2002). Less is known about the role of mitochondria at the post-synapse. Previous work demonstrated that the proper localization of mitochondria in cultured hippocampal neuron dendrites is also critical to support synapses. Specifically, increasing either mitochondrial content or mitochondrial activity in dendrites increased the number of spines and synapses (Li et al. 2004). In turn, synaptic activity can regulate the motility of mitochondria in dendrites, leading to an accumulation of mitochondria near recently active synapses (Li et al. 2004). Interestingly, the morphology and the distribution of mitochondria differs between axons and dendrites *in vivo* and *in vitro* (Chang, Honick, and Reynolds 2006; Lewis et al. 2018), and mitochondria in the two compartments are differentially modulated by synaptic activity in culture (Chang, Honick, and Reynolds 2006). However, whether the morphology and the distribution of mitochondria differs across hippocampal subregions and compartments is not known.

We previously reported that CA2 of the hippocampus, a subregion that is essential for social memory (Hitti and Siegelbaum 2014; Stevenson and Caldwell 2014; Dudek, Alexander, and Farris 2016), is enriched for transcripts related to mitochondrial function compared to neighboring subregions CA1, CA3, and Dentate gyrus, DG (Farris et al. 2019). In particular, we found that multiple components of the mitochondrial calcium uniporter (MCU) complex are enriched in CA2 (Farris et al. 2019). The MCU complex regulates calcium entry into the mitochondrial matrix (Baughman et al. 2011; De Stefani et al. 2011), which can induce morphological (Han et al. 2008) and metabolic changes (Wescott et al. 2019; Llorente-Folch et al. 2015) and regulate the transport and localization of mitochondria to active synapses (Wang and Schwarz 2009). Thus, enrichment of MCU in CA2 neurons may differentially influence mitochondrial morphology and localization to meet local energy demands.

In this study, we found that MCU expression was strikingly enriched in CA2 distal apical dendrites compared to proximal apical dendrites, an enrichment not seen in neighboring CA1 distal apical dendrites. MCU-enriched mitochondria in CA2 distal dendrites were more tubular in shape and significantly larger in size compared to mitochondria in CA2 proximal dendrites and CA1 dendrites, although mitochondria in CA1 distal dendrites were also larger than mitochondria in CA1 proximal dendrites. Distal dendrites in the hippocampus receive distinct inputs compared to proximal dendrites, demonstrating that mitochondria display cell- and circuit-specific heterogeneity in the hippocampus with a particularly unique subpopulation in CA2 distal dendrites. Overexpression of MCU in CA1 significantly increased mitochondrial mass and number throughout CA1 apical dendrites. However, this effect was preferentially seen in the proximal dendrites compared to the distal dendrites, suggesting an MCU-induced swap in mitochondrial localization within the dendrites of CA1. These data shine light on the molecular and morphological diversity of mitochondria across hippocampal cell types and circuits, and suggest that MCU enrichment in CA2 distal dendrites could be a mechanism for differentially regulating layer-specific synaptic energy demands, and thus CA2 circuit function in social memory.

## RESULTS

### MCU is enriched in CA2 distal dendrites and reveals morphological differences across proximal and distal dendrites

In order to assess the heterogeneity of MCU expression across hippocampal circuits, we immunostained hippocampal-containing sections from C57bl6/J mice for MCU. Consistent with our transcriptome study (Farris et al. 2019), MCU fluorescence was highly enriched in RGS14-positive CA2 cell bodies and dendrites compared to neighboring CA1 cell bodies and dendrites (Fig. 1A-D, two-way RM ANOVA, effect of subregion: F =309.8, P = 0.003; effect of layer: F = 24.5, P = 0.006; effect of subregion x layer: F = 13.44, P = 0.017). However, within CA2, MCU fluorescence was strikingly enriched in the distal apical dendrites in layer stratum lacunosum moleculare (SLM) compared to the proximal apical dendrites in layer stratum radiatum (SR, Fig.1D, Sidak’s multiple comparison test, CA2 SR vs SLM: P = 0.011). MCU fluorescence was not detected in interneurons or non-neuronal cells present in SR or SLM layers. The same level of MCU fluorescence enrichment in CA2 SLM was not seen in neighboring CA1 SLM (Fig. 1D, CA1 SR vs SLM: P = 0.473). The distal dendrites of CA2 (SLM) receive layer-specific input from entorhinal cortex layer II, whereas the proximal dendrites (SR) receive input from CA3. These data suggest that there is mitochondrial diversity across hippocampal cell types, and even within dendritic domains of the same cells which reflect distinct hippocampal circuits.

**Figure 1:**
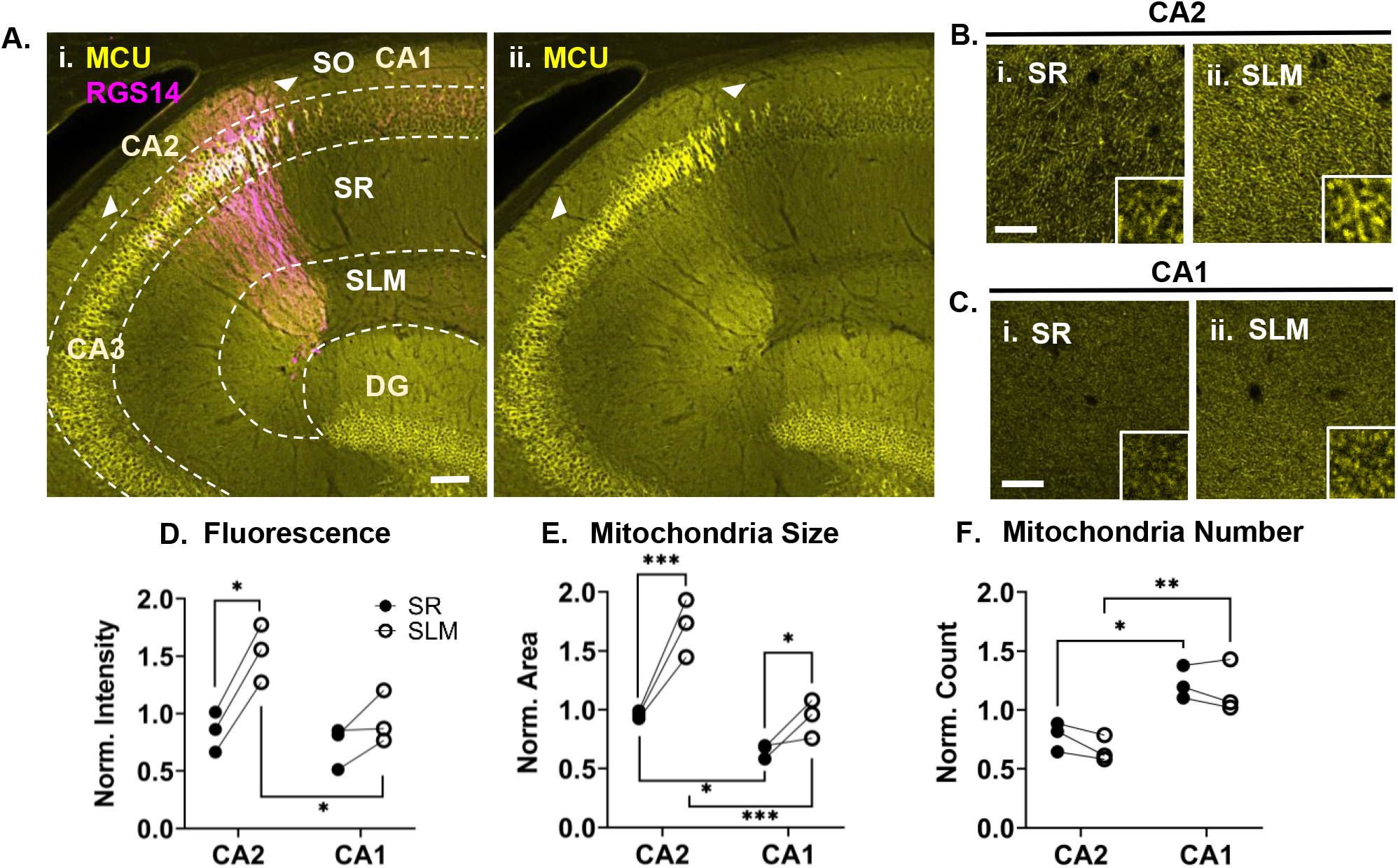
MCU-labeled mitochondria are preferentially enriched in CA2 distal dendrites and larger in size than neighboring mitochondria. **A.** (*i*.) 20X epifluorescence tile image of a wild-type hippocampus stained for MCU (yellow) and RGS14 (magenta) with the subregions and dendritic layers labeled. White arrowheads indicate the borders of CA2. (*ii*.) The MCU channel alone. **B.** Representative 40X confocal images of MCU staining in SR (*i*.) and SLM (*ii*.) of CA2 in a wild-type mouse. All representative 40X images are from a single Z section. Insets in **B** and **C** are a zoomed 10 X 10 μm crop of the larger image. **C.** Representative 40X confocal images of MCU staining in SR (*i*.) and SLM (*ii*.) of CA1 in a wild-type mouse. Images were processed identically to the representative images in **B** for comparison between CA1 and CA2. **D.** The mean MCU fluorescence of a representative 100 X 100 μm ROI from SR and SLM of CA1 and CA2. Data were normalized to the overall average. SR (closed circles) and SLM (open circles) of the same animal are paired with lines. Two-way RM ANOVA with Sidak’s post hoc; N = 3 animals; 4-5 hippocampal sections / animal. Overall effect of subregion and layer were significant (subregion: F = 309.8, P = 0.003; layer: F = 24.5, P = 0.006; interaction: F = 13.4; P = 0.017). **E.** The average area of MCU-labeled mitochondria in the same ROIs from **D.** Mitochondria were segmented as described in the methods. Data were normalized and plotted the same as in **D.** Two-way RM ANOVA with Sidak’s post hoc. Overall effect of subregion and layer on area were significant (subregion: F = 191.0, P = 0.005; layer: F = 18.0, p = 0.01, interaction: F = 62.1; P = 0.001). **F.** The average count of MCU-labeled mitochondria from the dataset in **E.** Two-way RM ANOVA with Sidak’s post hoc. Overall effect of subregion was significant (F = 515.8, P = 0.002). Overall effect of layer on count was significant (F = 12.4, p = 0.019); however, none of the post hoc comparisons between dendritic layers were significant. SO = Stratum Oriens; SR = Stratum Radiatum, SLM = Stratum Lacunosum Moleculare. Sidak’s post hoc: * = P < 0.05; ** = P < 0.01; *** = P < 0.001. Scale for A = 100 μm; B-C = 20 μm.

To quantify differences in mitochondrial morphology across the dendritic layers of CA1 and CA2, individual MCU-labeled mitochondria were segmented and quantified to obtain the average mitochondrial area and count for each dendritic layer (see methods). Supplemental Figure 1 includes the data and statistics for all three dendritic layers (stratum oriens (SO), SR, and SLM); however, we focused our analyses on layers SR and SLM, as less is known about CA2 basal dendrite connectivity and plasticity and thus results would be hard to interpret. We found that mitochondrial area varied by subregion and dendritic layer (two-way RM ANOVA, effect of subregion: F = 191.0, P = 0.005; effect of layer: F = 18.0; P = 0.01).The average MCU-labeled mitochondrial area was significantly greater in SLM compared to SR in CA2 and, to a lesser extent, in CA1 (Fig. 1E, Sidak’s multiple comparison test, CA2 SR vs SLM: P = 0.0005; CA1 SR vs SLM: P = 0.023). Across both dendritic layers, MCU-labeled mitochondrial area was significantly larger in CA2 than in CA1 (CA1 SR vs CA2 SR: P = 0.016; CA1 SLM vs CA2 SLM; P = 0.0004). Interestingly, the number of mitochondria was significantly less in CA2 than in CA1 for all dendritic layers (Fig 1F; two-way RM ANOVA, effect of subregion: F = 515.8, P = 0.002; Sidek’s multiple comparison test, CA1 vs CA2 SR: P = 0.018; SLM: P = 0.01). Although there was an overall effect of layer (F = 12.4, P = 0.019), mitochondrial number was not significantly different between SR and SLM of CA1 or CA2 (CA1 SR vs SLM: P = 0.999, CA2 SR vs SLM: P = 0.742). Thus, mitochondria appear to be generally larger in the distal dendrites of both CA1 and CA2 compared to their proximal dendrites. However, MCU-enriched mitochondria in CA2 dendrites are larger in size and fewer in number compared to mitochondria in CA1 dendrites. Further, these effects are most evident in CA2 distal dendrites, indicating that CA2 distal dendrites contain a unique subpopulation of mitochondria.

### COX4 labels a broader but overlapping population of mitochondria than MCU

In addition to MCU, we stained for mitochondria using an antibody against cytochrome c oxidase subunit 4 isoform 1 (COX4), a component of the enzyme complex involved in the terminal step of the mitochondrial electron transport chain. Similar to MCU, COX4 staining was enriched in CA2 cell bodies compared to neighboring CA1 cell bodies (Fig. 2A). However, COX4 dendritic staining was more variable and less striking than MCU dendritic staining and thus required a greater number of mice to assess dendritic mitochondrial morphology. This blunted COX4 dendritic expression profile compared to MCU may be due to COX4 expression in non-pyramidal cells within the dendritic laminae (Fig. 2A(ii) yellow arrows). Comparing COX4 labeling in the dendritic layers of CA2 and CA1 (Fig. 2BC) revealed small, but significant differences in COX4 fluorescence (Fig. 2D, two-way RM ANOVA, effect of layer: F = 13.4, P = 0.01) and COX4-labeled mitochondria size (Fig. 2E, two-way RM ANOVA, effect of layer: F = 15.5, P = 0.008). COX4 fluorescence was increased in SLM compared to SR in both CA2 and CA1 dendrites (Fig. 2D, Sidak’s multiple comparison test, CA2 SR vs SLM: P = 0.0006; CA1 SR vs SLM: P = 0.092). Similarly, COX4-labeled mitochondrial area was increased in SLM compared to SR in both CA2 and CA1 (Fig. 2E, Sidak’s multiple comparison test, CA2 SR vs SLM: P = 0.002; CA1 SR vs SLM: P = 0.003). Unlike with MCU-labeled mitochondria, there was no statistical difference in COX4 fluorescence or COX4-labeled mitochondrial area between CA2 and CA1 dendrites (Fig. 2DE, two-way RM ANOVA, effect of subregion: P > 0.05 for fluorescence and area). There was also no significant difference in the number of COX4-labeled mitochondria in CA2 versus CA1, or across dendritic layers (Fig. 2F, two-way RM ANOVA, effect of subregion and layer: P > 0.05). In sum, COX4-labeled mitochondria show no significant differences across subregions CA1 and CA2, but show consistent increases in fluorescence and mitochondrial area in distal dendrites compared to proximal dendrites. Thus, the selective enrichment of MCU in the distal dendrites of CA2 compared to CA1 appears to be unique, suggesting the mitochondria there are molecularly distinct.

**Figure 2:**
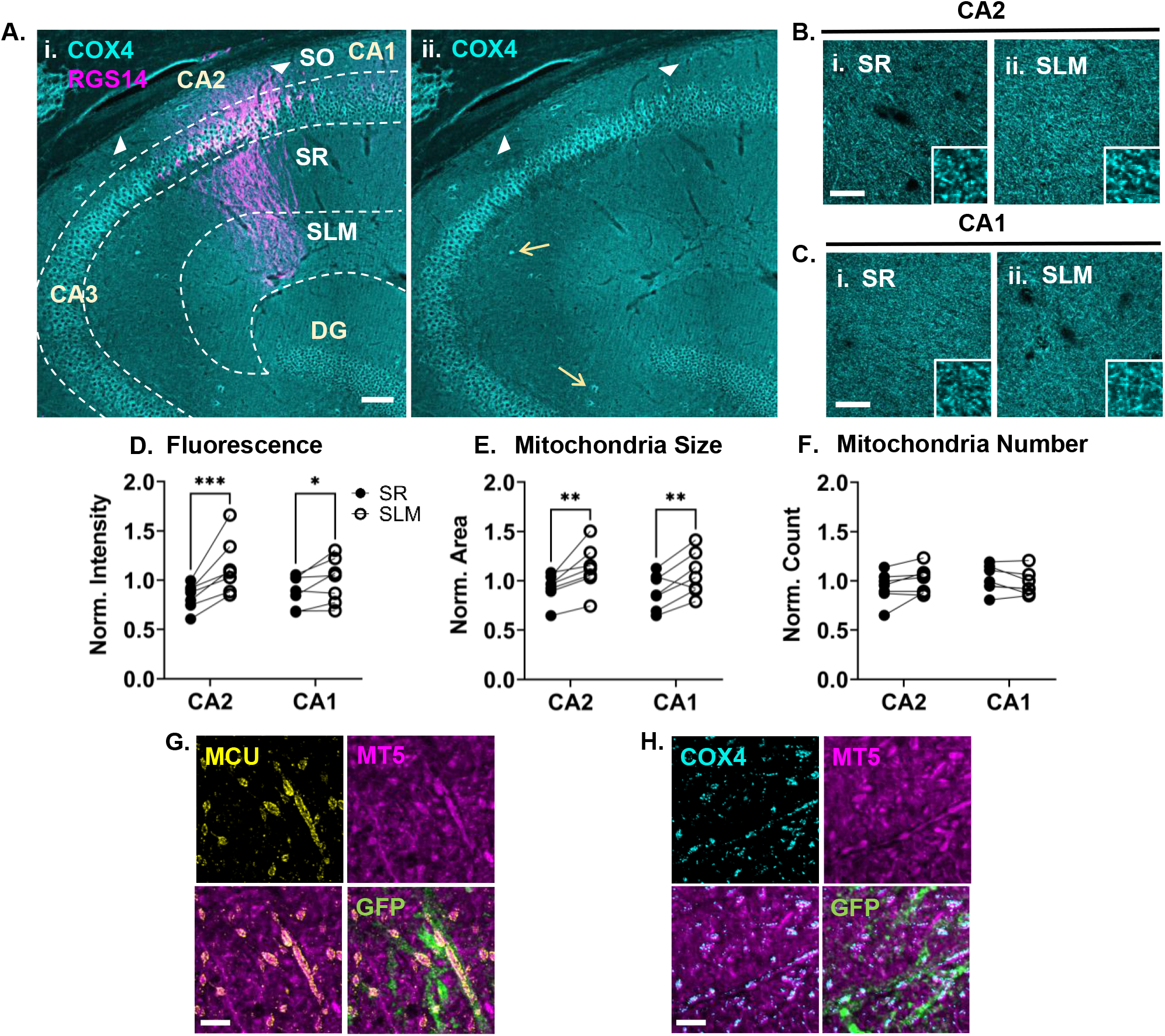
COX4 labeling is similarly enriched in CA2 and CA1 distal dendrites. **A.** (*i*.) 20X epifluorescence tile image of a wild-type hippocampus stained for COX4 (cyan) and RGS14 (magenta) with the subregions and dendritic layers labeled. White arrowheads indicate the borders of CA2. (*ii*.) The COX4 channel alone. Yellow arrows show examples of COX4 labeling in non-pyramidal cells in the neuropil. **B.** Representative 40X confocal images of COX4 staining in SR (*i*.) and SLM (*ii*.) of CA2 in a wild-type mouse. Insets in F and G are a zoomed 10 X 10 μm crop of the larger image. **C.** Representative 40X confocal images of COX4 staining in SR (*i*.) and SLM (*ii*.) of CA1 in a wild-type mouse. Images in F and G were processed identically to preserve differences between subregions and dendritic layers. **D.** COX4 mean fluorescence of a 100 x 100 urn ROI from SR and SLM of CA1 and CA2 normalized to the overall cohort average. Lines pair SR and SLM of the same animal. Two-way RM ANOVA with Sidak’s post hoc; N = 7 animals from two cohorts. Overall effect of subregion was not significant for any metric (p > 0.05), thus subregion comparisons are not shown. Overall effect of layer was significant for fluorescence (F = 13.4, p = 0.011). **E.** Mean area of COX4-labeled mitochondria in the same ROIs in **D.** Mitochondria were segmented the same as in Figure 1 and data were normalized to the overall cohort average. Two-way RM ANOVA with Sidak’s post hoc. Overall effect of layer on area was significant (F = 15.5, p = 0.008). **F.** Mean count of COX4-labeled mitochondria from the same dataset as **E.** Two-way RM ANOVA with Sidak’s post hoc. The overall effect of layer and subregion on mitochondria count were not significant (P > 0.05). **G.** Representative ProExM images of MCU (yellow) and MT-5 (magenta) in CA2 SLM. Bottom panels show the overlay of both channels (left) and an overlay with GFP+ CA2 distal dendrites (right). **H.** Representative ProExM images of COX4 staining (yellow) and MT-5 staining (magenta) in CA2 SLM. An ROI was chosen which had a similar amount of MT-5 and GFP staining as in **G** for comparison. Sidak’s post hoc: * = P < 0.05; ** = P < 0.01; *** = P < 0.001. Scale for A = 100 μm; B-C = 20 μm; G-H = 3 μm (ExM adj.)

To test the possibility that MCU is labeling a subpopulation of COX4-labeled mitochondria, we acquired super resolution images of MCU- (Fig. 2G) and COX4- (Fig. 2H) labeled mitochondria in CA2 SLM alongside a novel pan-mitochondrial matrix marker MT-5 (Kalyuzhny et al. 2021). We used protein retention expansion microscopy (ProExM) to visualize densely packed mitochondria (Campbell et al. 2021). In order to compare MCU and COX4 labeling across similar mitochondrial populations, we used ProExM images with similar MT-5 and CA2 SLM dendrite content. Although MCU and COX4 both labeled the inner mitochondrial membrane surrounding MT-5 labeled mitochondrial matrix, MCU-labeled mitochondria were qualitatively larger and less numerous than COX4-labeled mitochondria. This suggests that MCU may label a subset of the mitochondrial population labeled with COX4, perhaps only mitochondria in CA2 dendrites versus mitochondria from multiple cell populations within the neuropil.

### Genetically labeled mitochondria in CA2 distal dendrites are increased in size relative to the proximal dendrites

To confirm the difference in mitochondrial mass between the proximal and distal dendrites of CA2, we used a genetic strategy to sparsely label CA2 neurons with GFP-tagged mitochondria. We crossed Mitotag mice (Fecher et al. 2019) to a tamoxifen-inducible CA2-specific cre line (Amigo2-icreERT2, Alexander et al. 2018), which allowed for mitochondria to be resolved in individual CA2 neuron dendrites (Fig. 3A). Consistent with our MCU immunolabeling, mitochondria size varied by CA2 dendrite layer with smaller, punctate mitochondria in SO compared to enlarged, tubular mitochondria in SLM, with SR mitochondria sized in between. Similar results were qualitatively seen in scanning electron micrographs (SEM, Fig. 3B). The median area of Mitotag-labeled mitochondria in SLM was statistically greater than Mitotag-labeled mitochondria in both SO and SR (Fig. 3C, RM One Way ANOVA, F=23.47, P=0.0014, Tukey’s multiple comparison test, SO vs. SLM: P=0.009; SR vs. SLM: P=0.007). The ratio of median mitochondrial areas for SLM to SR was consistently above 1.0 across mice (Fig. 3E, average SLM:SR ratio =1.24 ± 0.04 SEM, N=5 mice) and the relative frequency curve for individual mitochondrial areas in SLM was shifted to the right compared to individual mitochondrial areas in SO and SR (Fig. 3E). Collectively, these findings indicate that layer-specific differences in mitochondrial morphology are not due to an apparent increase in mitochondrial size due to enriched MCU or COX4 expression.

**Figure 3:**
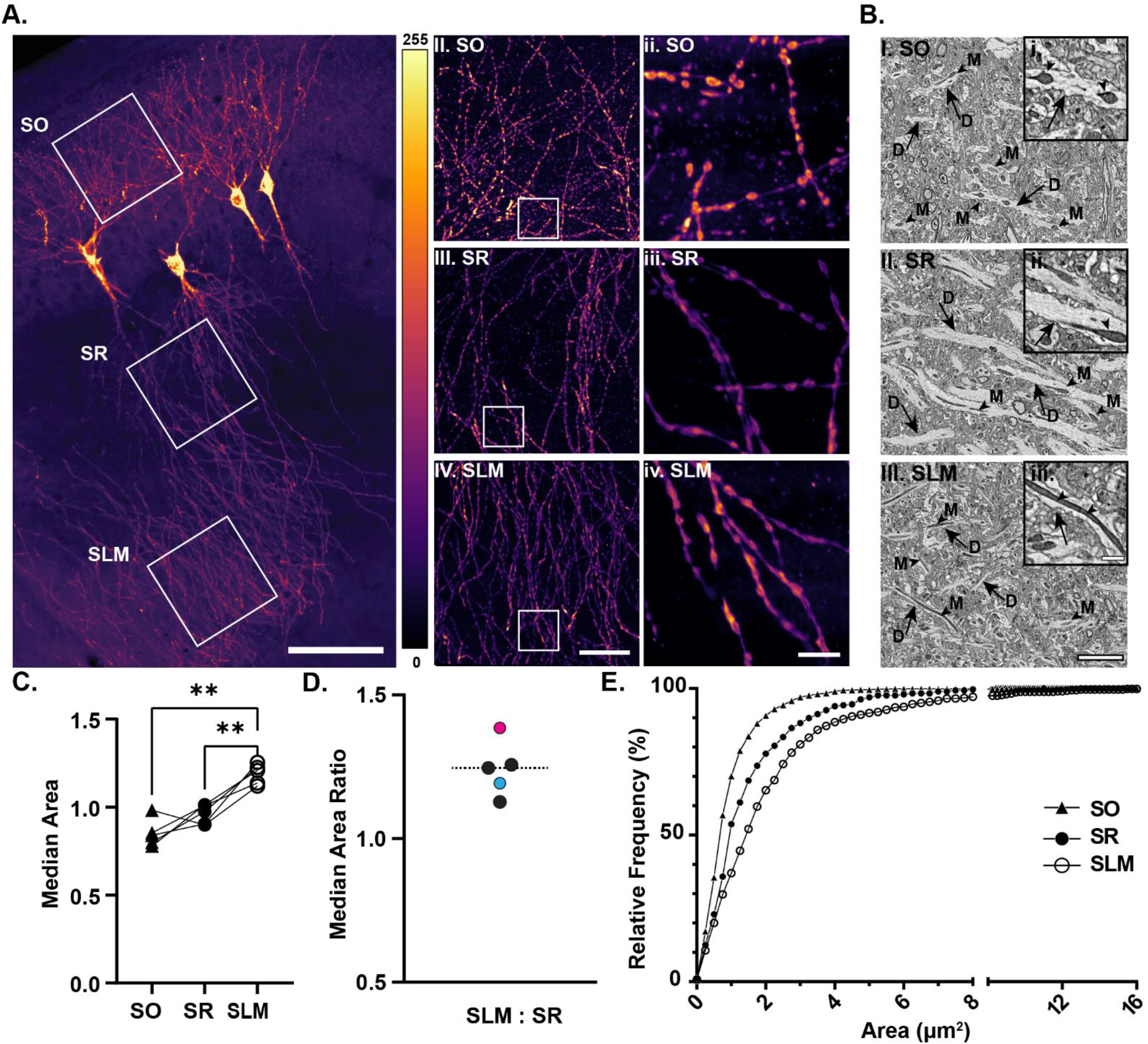
Super resolution imaging and electron microscopy confirm CA2 layer-specific differences in dendritic mitochondrial morphology. **A.** (I) Representative 20X confocal image of sparse CA2 neuron Mitotag labeling of mitochondria. ROIs (white box) within the 20X image indicate the selection for6OX (II) SO, (III) SR, and (IV) SLM representative images. ROI (white box) within 6OX images is representative of 10×10 μm^2^ 4X super-resolution images of Mitotag labeled mitochondria in individual (ii) SO, (iii) SR, and (iv) SLM dendritic branches. **B.** Representative electron micrographs of (i) SO, (ii), SR, and (iii) SLM dendrites. Examples of mitochondria (arrowheads) within dendrite (arrows) cross sections are indicated. **C.** Median Mitotag-labeled mitochondrial area in CA2 SO, SR, and SLM dendritic layers. Data are normalized to the average area per mouse with lines connecting SO, SR, and SLM per mouse. Mitochondrial size was significantly increased within the SLM layers compared to both SO and SR in the CA2 (RM One Way ANOVA: F=23.47, P=0.0014, Tukey’s multiple comparison test; N=5 mice; 4-7 hippocampus sections/animal). **D.** Ratio of the median mitochondrial area between SLM and SR layers in C. Color-filled data points represent the mice selected for representative images (A: blue; E: pink). **E.** Percent relative frequency plot of individual mitochondrial areas from CA2 SO, SR, and SLM from one representative section from the mouse labeled in pink in D. SO = Stratum Oriens, SR = Stratum Radiatum, SLM = Stratum Lacunosum Moleculare. Tukey’s post hoc: * = P <0.05; ** = P <0.01. Scale = (AI) 100μm, (AIV) 25μm, (Aiv, BIII) 5μm, and (Biii) 1 μm.

### MCU overexpression influences dendritic mitochondrial mass and layer-specific localization

Although the exact role of MCU in regulating mitochondrial function is still being explored, it has been shown that calcium influx into the mitochondrial matrix stimulates ATP production via the TCA cycle (Wescott et al. 2019; Llorente-Folch et al. 2015), causes morphological changes by inducing mitochondrial fragmentation via fission (Han et al. 2008), and regulates the transport of mitochondria and the localization of mitochondria at active synapses (Wang and Schwarz 2009). However, whether cell-specific differences in MCU expression contribute to differences in mitochondrial morphology or localization is unknown.

Based on our findings that MCU expression correlates with increased mitochondrial size in the distal dendrites of CA2, we next asked whether increasing MCU expression can increase mitochondrial size or influence mitochondrial localization in the dendrites of neighboring CA1. To test this, we used adeno-associated viral vectors (AAVs) to overexpress MCU or GFP in CA1 neurons, which normally have 4-fold less *Mcu* expression than CA2 neurons (Farris et al. 2019). CA1 was stereotaxically targeted in wild-type mice to infuse either AAV-MCU (AAV9-hSYN1-mMCU-2A-GFP-WPRE) or AAV-GFP (AAV9-hSYN1-eGFP) as a control. We then immunostained for MCU and GFP and quantified the mean MCU fluorescence and the size and number of MCU-labeled mitochondria in SO, SR and SLM of CA1. MCU expression was about ~4.5-fold greater overall in the AAV-MCU treated mice than the control AAV-GFP treated mice (Fig. 4AB). MCU overexpression in CA1 resulted in larger (two-way RM ANOVA, effect of AAV: F = 124.4, P = < 0.0001) and more numerous (two-way RM ANOVA, effect of AAV: F = 12.03, P = < 0.007) MCU-labeled mitochondria in CA1 dendrites compared to GFP control mice (Fig. 4C). Fig. 4C shows the effect of AAV treatment on mitochondrial area versus mitochondrial number for AAV-MCU (red diamonds) and AAV-GFP (blue circles) treated mice. In AAV-MCU treated mice, there was an increase in both mitochondrial size and number compared to AAV-GFP treated mice, shifting the data towards the upper quadrant of the plot (Fig. 4C). This confirms that MCU overexpression is sufficient to increase mitochondrial content in CA1 dendrites. However, the effect of MCU overexpression was greater in CA1 SR compared to SLM for MCU fluorescence (Fig. 4D, Sidak multiple comparisons; P = 0.002) and mitochondrial size (Fig. 4E; Sidak multiple comparisons, P = 0.005). This can also be seen in the correlation plot in Fig. 4C, where in AAV-MCU treated mice there are larger and more numerous mitochondria in SR compared to SLM, whereas in control mice, it is the opposite. This indicates that, in contrast to CA2 dendrites where MCU expression is enriched in mitochondria in distal dendrites, MCU overexpression in CA1 preferentially influences mitochondrial size and localization in proximal dendrites compared with distal dendrites. As with CA2, proximal dendrites in CA1 receive layer specific input from area CA3, but in contrast to CA2, CA1 distal dendrites receive input from entorhinal cortex layer III. Again, suggesting that distinct hippocampal circuits are differentially regulated by diverse pools of mitochondria. Mitochondria labeled with MT-5 also appear to be qualitatively larger and fewer in number in CA1 SR of AAV-MCU treated mice compared to AAV-GFP treated mice (Fig. 4G). In particular, long tubular MT-5 labeled mitochondria observed in AAV-MCU treated mice were generally colocalized with MCU (white arrowhead in Fig. 4G(ii.)).

**Figure 4:**
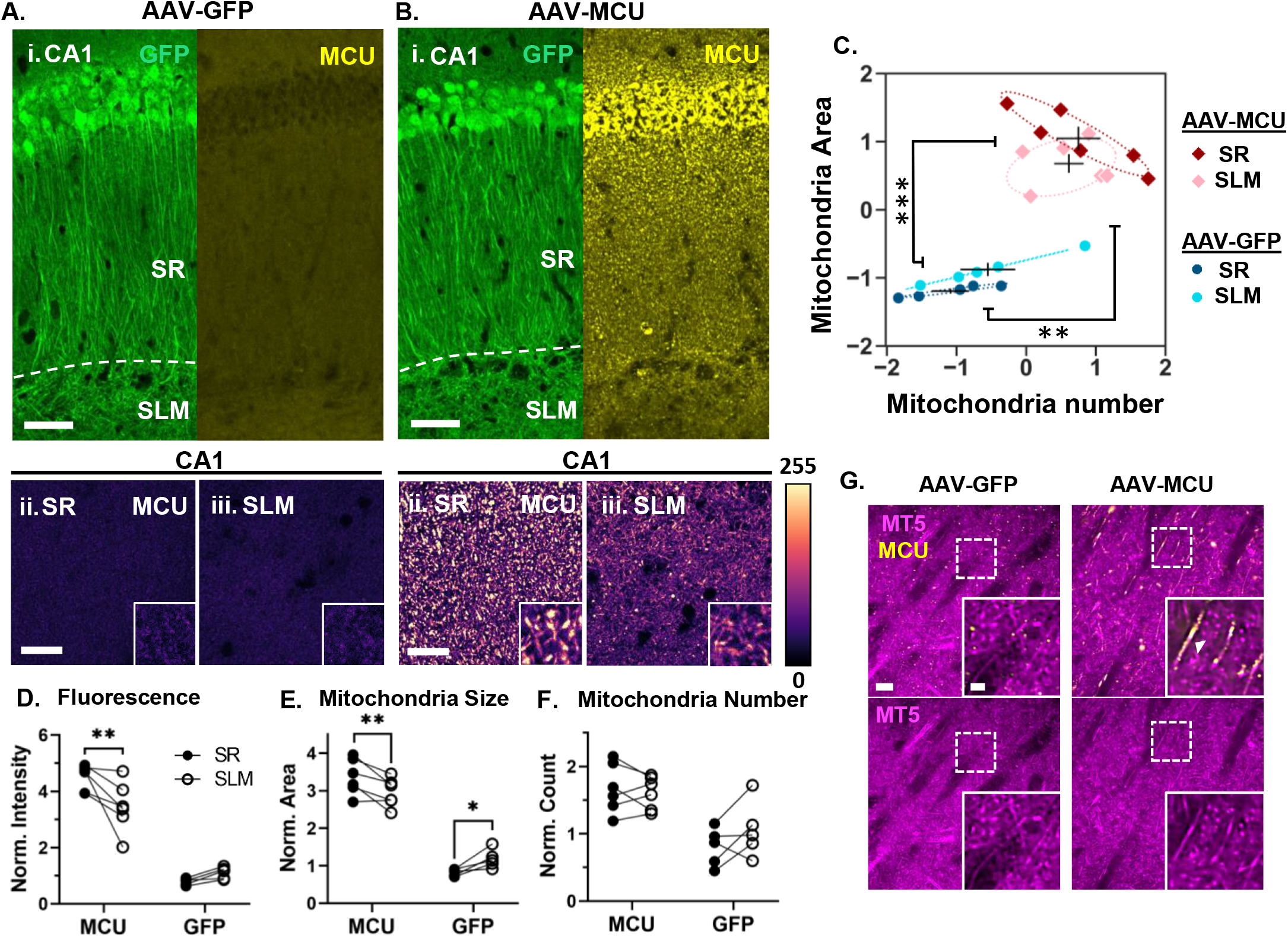
CA1 MCU overexpression preferentially increases mitochondrial mass and localization in proximal dendrites. **A.** (i) Representative 20X epifluorescence image of CA1 stained for GFP (left) and MCU (right) in a AAV-GFP treated control animal. Bottom: Representative 40X confocal images of MCU staining in SR (i.) and SLM (ii.) of CA1 in a AAV-GFP treated mouse. Inset is a zoomed 10 X 10 μm crop. **B.** (i) Representative 20X epifluorescence image of CA1 stained for GFP (left) and MCU (right) in an AAV-MCU treated mouse. Images in A and B were processed the same to maintain relative difference between AAV-MCU and AAV-GFP mice. Bottom: Representative 40X confocal images of MCU staining in SR (i.) and SLM (ii.) of CA1 in an AAV-MCU treated mouse. **C.** Correlation plot of mitochondrial area versus mitochondrial number in SR (dark shades) and SLM (light shades) of AAV-GFP (blue circles) and AAV-MCU animals (red diamonds). Average Z score is plotted for each animal, cross-hairs show the mean and standard error. An ellipse was fit to the data for each group (dotted line). Stats bars show the overall effect of AAV treatment from a two-way RM ANOVA. AAV-MCU treatment significantly increased mitochondria area and number relative to AAV-GFP (Area: F = 124.4, P <0 0001; Number: F = 12.03, P <0.007). N = 6 AAV-MCU mice and 5 AAV-GFP controls. **D.** Mean MCU fluorescence of a 100 X 100 μm ROI from SR and SLM of the AAV-MCU and AAV-GFP mice from **C** normalized to GFP control average. Two-way RM ANOVA with Sidak’s post hoc comparing dendritic layers. Overall effect of AAV treatment and layer were significant (Treatment: F = 113.4, P = <0.0001; Layer: F = 4.81, P = 0.021). See supplemental Table 1 for post hoc comparisons. **E.** MCU-labeled mitochondrial area in SR and SLM of the AAV-MCU and AAV-GFP mice normalized to GFP control average. Two-way RM ANOVA with Sidak’s post hoc. Overall effect of layer was significant (F = 3.84, P = 0.041). **F.** Mitochondrial number in SR and SLM of the AAV-mice and AAV-GFP mice normalized to GFP control average. Two-way RM ANOVA with Sidak’s post hoc. Overall effect of layer was significant (F = 4.60, P = 0.024). **G.** Representative ProExM images of MT-5 (magenta) and MCU labeling (yellow) in CA1 SR of an AAV-GFP (i) and AAV-MCU (ii) mouse. Dashed box shows the location of the zoomed image in the inset. White arrow shows an example of a large MT-5 and MCU+ mitochondria. Bottom: Same images as above, but only the MT-5 channel. SR = Stratum Radiatum; SLM = Stratum Lacunosum Moleculare. Sidak’s post hoc: * = P < 0.05; ** = P < 0.01. Scale for A-B(i.) = 50 μm; Scale for A-B(ii.-iii.) = 20 μm; Scale for G = 3 μm, inset = 1 μm (ExM adj.)

## DISCUSSION

The goal of this study was to investigate mitochondrial heterogeneity in the mouse hippocampus. Focusing on the enrichment of MCU in area CA2, we unexpectedly uncovered that MCU expression is preferentially enriched in CA2 distal apical dendrites, precisely where CA2 neurons receive layer-specific input from entorhinal cortex II, a circuit recently shown to carry social information to CA2 (Lopez-Rojas et al. 2022; Dang et al. 2022). This striking enrichment of MCU in CA2 distal dendrites compared to CA2 proximal dendrites and neighboring CA1 dendrites is not seen with another mitochondrial marker, COX4, suggesting that mitochondria in CA2 distal dendrites have a unique molecular expression profile. We further found that MCU-enriched mitochondria in CA2 distal dendrites are larger and more tubular compared with mitochondria in CA2 proximal dendrites and CA1 apical dendrites. The increased mitochondrial size in the distal dendrites of CA2 was not limited to MCU-labeled mitochondria, as this was seen in COX4-labeled mitochondria, as well as genetically labeled mitochondria and mitochondria in electron micrographs. Studies in cultured hippocampal neurons have demonstrated that elongated mitochondria in axons can take up more calcium, suggesting a direct connection between mitochondrial morphology and function (Lewis et al. 2018). This is supported by our finding that MCU overexpression in CA1 increases mitochondrial size and number preferentially in the proximal dendrites of CA1, which are more efficient at triggering somatic excitatory postsynaptic currents (EPSCs, Srinivas et al. 2017) and are more plastic than CA1 distal dendrites (Sajikumar and Korte 2011), which collectively should produce more calcium. While further studies are required to establish whether MCU expression directly regulates mitochondrial mass and function in CA2 distal dendrites, our results revealed unexpected mitochondrial diversity across hippocampal subregions and compartments. In particular, we identified a molecularly and structurally distinct subpopulation of mitochondria in CA2 distal dendrites that could influence circuit specific forms of plasticity and behavior.

### Evidence for a distinct subpopulation of mitochondria in CA2 distal dendrites

Several lines of evidence support the finding that CA2 distal dendrites harbor larger and more tubular mitochondria compared to CA2 proximal dendrites and CA1 apical dendrites. Chiefly, MCU-labeled mitochondria are visibly and quantitatively larger in CA2 distal dendrites compared to CA2 proximal dendrites and CA1 apical dendrites. Quantification of GFP tagged mitochondria in sparsely labeled CA2 dendrites closely mirrored the results from MCU-labeled mitochondria. Moreover, we provided electron microscopic evidence for differentially sized and shaped mitochondria in each dendritic layer consistent with results from the other methods. However, while COX4 labeled mitochondria showed similar, albeit smaller, differences in fluorescence and mitochondrial size between CA2 proximal and distal dendrites, we did not detect COX4 labeled mitochondrial differences between CA1 and CA2, as strikingly seen with MCU-labeled mitochondria. We noted that COX4 labeled mitochondria appear to be more numerous than MCU-labeled mitochondria across comparative populations, and that COX4 labels mitochondria in non-pyramidal cells within the neuropil, whereas MCU only labels mitochondria in pyramidal neurons. Thus, COX4-labeling likely includes a broader population of mitochondria beyond mitochondria in CA2 or CA1 dendrites. MCU may be labeling a subset of mitochondria that partially overlap with COX4 labeled mitochondria. However, we cannot exclude the possibility that the quantitative differences between MCU- and COX4-labeled mitochondria are due to differences in the quality of antibody labeling. Further studies are required to incisively elucidate whether MCU-enriched mitochondria in CA2 distal dendrites are molecularly distinct, or whether they just appear as such due to their greater mitochondrial content.

### Potential functional roles for a distinct subpopulation of mitochondria in CA2 distal dendrites

There are at least two possible reasons for why mitochondria are different in CA2 distal apical dendrites. Firstly, the CA regions receive dendritic layer-specific inputs which have different efficacies for generating somatic responses. Inputs that are more proximal to the soma typically cause a larger somatic EPSC; however, ECII inputs onto the distal dendrites of CA2 drive an unusually robust response in CA2 neurons, 5 to 6-fold larger EPSCs than those evoked from ECIII inputs onto CA1 neurons (Srinivas et al. 2017, but see also Sun et al. 2021). This enhanced efficacy is the result of unique synaptic and dendritic properties in CA2 SLM (Srinivas et al. 2017; Sun et al. 2014), including almost 3 times as many glutamatergic synapses in CA2 SLM than in CA1 SLM (Srinivas et al. 2017), although there is heterogeneity in CA2 dendritic SLM morphology (Helton et al. 2019). Moreover, there is an asymmetrical gradient of innervation from ECII, with the lateral entorhinal cortex II (LECII) axons covering a larger area more distal from the soma than the medial entorhinal cortex II axons (MECII) (Steward, 1976). This is consistent with the asymmetric pattern of MCU-labeled mitochondrial morphology in CA2 dendrites. Notably, input from the LECII, but not the MECII, carries social information relevant to social recognition memory (Lopez-Rojas et al. 2022). Taken together, CA2 SLM may have higher expression of MCU and greater mitochondrial mass than CA1 SLM in order to metabolically support more numerous and more active synapses. In the cerebellum, higher expression of MCU in granule cells compared to Purkinje cells and astrocytes correlated with greater mitochondrial calcium uptake and buffering capacity (Fecher et al. 2019), which is associated with an increase in ATP production (Wescott et al. 2019). Future studies will be needed to expand on whether there are layer specific differences in metabolic demand that depend on MCU.

Secondly, the proximal and distal circuits within CA2 dendrites also have distinct plasticity profiles. Synapses from CA3 onto CA2 SR are known to be resistant to long-term potentiation (LTP, Zhao et al. 2007; Simons et al. 2009), while synapses in CA2 SLM from the entorhinal cortex readily express LTP (Chevaleyre and Siegelbaum 2010). As a result, the more plastic synapses in CA2 SLM may have a higher metabolic demand than the less plastic synapses in CA2 SR. It is known that mitochondria are actively recruited to the presynapse, where it is thought that they locally regulate ATP and cytoplasmic calcium to support synaptic transmission (Chang, Honick, and Reynolds 2006). In axons, MICU3, a brain-specific regulator of MCU, is necessary for activity-dependent mitochondrial calcium uptake and ATP synthesis to fuel synaptic transmission (Ashrafi et al. 2020). At the post-synapse, repeated stimulation of cultured hippocampal neurons induces mitochondria to invade enlarged spines in an NMDA-dependent manner (Li et al. 2004), which in theory could support a plasticity-induced increase in energetic demand. Studies have further shown that dendritic mitochondria in cultured neurons serve as a local energy source to fuel local translation-dependent changes at synapses during plasticity (Rangaraju et al. 2019). Similarly, the tubular, MCU-enriched mitochondria in the distal dendrites of CA2 could support the increased propensity for plasticity at these synapses. Given the role of MCU in calcium buffering and homeostasis, it is also possible that MCU plays a more direct role in LTP by regulating cytoplasmic calcium levels. We have previously shown that pharmacological blockade of mitochondrial calcium influx through MCU with Ru360 results in changes to the plasticity-resistant profile of CA2 SR synapses (Farris et al. 2019). However, it remains to be tested how MCU-enriched CA2 SLM synapses respond to MCU blockade or MCU knockout. Other studies have shown that MCU overexpression can increase mitochondrial calcium uptake after NMDA receptor activation (Qiu et al. 2013). In addition, mitochondrial fission, which is regulated by calcium influx into mitochondria, is required for structural LTP in CA1 (Divakaruni et al. 2018). However, the exact role MCU might be playing in regulating the plasticity at CA2 SLM synapses is an open question.

### MCU as a potential regulator linking local synaptic activity to mitochondrial function

Overexpression of MCU in CA1 increased dendritic mitochondrial size and influenced layer-specific distribution. Specifically, we found that MCU overexpression caused a relatively greater increase in MCU-labeled mitochondrial size and number in proximal dendrites (SR) versus the distal dendrites (SLM) relative to control GFP mice. This may be explained by the fact that synapses in CA1 SR are more efficient at triggering action potentials, and therefore more metabolically demanding than synapses in CA1 SLM (Chevaleyre and Siegelbaum, 2010). We hypothesize that MCU-enriched mitochondria drive localization to dendritic domains with greater calcium influx and/or metabolic demand in order to match mitochondrial form and function to energy need. Contrary to this hypothesis, previous studies have shown that calcium influx into mitochondria leads to mitochondrial fission via phosphorylation of dynamin-related protein 1 (DRP1), resulting in more fragmented mitochondria (Zheng et al. 2017; Zhao et al. 2015; Kedra et al. 2021). However, the vast majority of these studies have been in disease models or in culture, and not much is known about the role of MCU in regulating mitochondrial mass in intact adult neuronal circuits as described in the present study. There is evidence that the balance of mitochondrial fission and fusion is regulated by activity in neurons (Li et al. 2004). Interestingly, age related changes in mitochondrial size have been reported in CA1 axons of adult rats compared to P15 rats in response to theta-burst stimulation (TBS), with TBS resulting in an increase in mitochondrial volume in adults and a decrease in mitochondrial volume at P15 (Smith et al. 2016). In addition, studies using isolated mitochondria from mouse embryonic fibroblasts have shown that increased oxidative phosphorylation stimulates mitochondrial fusion and elongation (Mishra et al. 2014). Thus, it is possible that MCU overexpression increases oxidative phosphorylation, which in turn increases mitochondrial mass. Taken together, the morphology and localization of mitochondria can vary dynamically across different cellular compartments based on specific local needs. We showed that MCU is a key regulator of, or contributor to, mitochondrial diversity in dendrites, specifically with regards to mitochondrial mass and asymmetric distribution. Local regulation of mitochondria via MCU could be an additional tool that neurons use to shape circuit-specific functions that drive behavior. Thus, differential expression of MCU across circuits in CA2 dendrites may contribute to CA2’s unique ability to encode social memory.

## METHODS

### Animals

The analysis of wild type MCU- and COX4-labeled mitochondria was done on adult Amigo2-EGFP reporter mice (RRID:MMRRC 033018-UCD) of both sexes, which were bred for at least 10 generations onto a C57BL/6J background. To generate inducible, CA2-specific Mitotag mice, Amigo2-iCreERT2 mice (Alexander et al. 2018) were crossed with homozygous Mitotag mice (Rosa26-CAG-LSL-GFP-OMM; Jax #: 032675; Fecher et al. 2019). Heterozygous Mitotag^fl/WT^; Amigo2-iCre^+/-^ male and female mice were intraperitoneally injected with Tamoxifen (100-150 mg/kg) over five consecutive days between postnatal (P) ages P26-61 and euthanized 4-12 days post injection for experiments in Figure 3. Adult male and female Ai14 TdTomato reporter mice on a C57BL/6J background (Jax #: 007914; Madisen et al. 2010) were used (in the absence of cre) for the AAV infusion experiments in Figure 4. Mice were group-housed under a 12:12 light/dark cycle with access to food and water ad libitum. All procedures were approved by the Animal Care and Use Committee of Virginia Tech.

### AAV stereotaxic infusions

Prior to surgery, mice received an intraperitoneal (IP) injection of ketamine/dexdomitor cocktail (ket 100mg/kg; dex 0.5mg/kg) for anesthesia. Mice were put on an absorbent pad and eyes covered with ophthalmic lubricant before being positioned into a stereotaxic apparatus with ear bars. The hair around the incision site was trimmed with electric clippers, and the incision area cleaned with 3 cycles of alternating betadine and 70% alcohol. After clearing the skull, a drill was used to make a unilateral burr hole targeting CA1 of the hippocampus (−2.1 AP; −1.4 ML; −1.4DV). Using a glass pipette attached to a syringe pump, 0.1-0.3μl of AAV-MCU (AAV9-hSYN-mMCU-2A-GFP-WPRE, Vector Biolabs, AAV-254662) or AAV-GFP (AAV9-hSYN1-eGFP, Vector Biolabs, VB1107) was infused into the hippocampus at a flow rate of 100 nl/min. After five minutes, the glass pipette was slowly retracted at a rate of 0.5 mm/min and the incision was closed with surgical glue. Mice were given a subcutaneous injection of buprenorphine (0.05 - 0.5 mg/kg) for analgesia and an IP injection of antisedan (atipamezole, 1 mg/kg) and allowed to recover on a heating pad until ambulatory. Two weeks after infusion, mice were given an overdose of sodium pentobarbital (150 mg/kg) and transcardially perfused with 4% paraformaldehyde then processed for immunohistochemistry as described below.

Due to the potential for MCU overexpression to cause cell death (Granatiero et al. 2019), we immunostained AAV treated sections for MAP2 and GFAP. We found no evidence of cell death or gliosis in CA1 of the AAV-MCU or AAV-GFP infused mice as was seen in mouse cortical neurons, but not zebrafish retina cone photoreceptors (Hutto et al. 2020). We presume this is an indication that different cell types have various capacities for MCU expression and/or mitochondrial calcium influx based intrinsic or extrinsic factors related to calcium signaling.

### Immunofluorescence

Mice were anesthetized with 150 mg/kg sodium pentobarbital and transcardially perfused with 15 ml of ice-cold 4% paraformaldehyde. Brains were dissected and post-fixed for at least 24 hours before sectioning 40-50 μm thick sections in the horizontal plane on a vibratome (Leica VT1000S). All brain sections immunostained with MCU or COX4 underwent antigen retrieval by boiling free floating sections for 3 min in 1ml of nanopure water, followed by a permeabilization step in 0.03% Triton-100x for 15 min at room temperature. All sections were then washed in PBS and blocked for 30-60 minutes in 5% Normal Goat Serum (NGS)/0.03% Triton-100x. Sections were incubated overnight (18-24 hours) with primary antibodies: rabbit-anti-MCU (1:500, Novus NBPS-76948), rabbit-anti-COX4 (1:500, SySy 298 003), chicken-anti-GFP antibody (1:500, Abcam ab13970), or a chicken-anti-RFP antibody (1:500, SySy 409-006) for any tissue containing the respective reporter. In some cases where CA2 was not labeled genetically, a mouse-anti-RGS14 antibody (1:500, NeuroMab 75-170), rabbit-anti-NECAB2 (1:500, Novus NBP2-84002), or rabbit-anti-PCP4 (1:1000, Invitrogen PA5-52209) was used to label CA2. After several rinses in PBS-T (0.03% Triton-100x), sections were given a quick second block (10-20 minutes) before incubating in secondary antibodies for 2 hours at room temperature. The following secondary antibodies were diluted 1:500 in blocking solution: Invitrogen goat-anti-mouse-488 A11029, goat-anti-chicken-488 A11039, goat-anti-rabbit-546 A11035, or Sigma goat-anti-mouse-633 SAB4600141. Prior to imaging, sections were washed in PBS-T and mounted under Prolong gold fluorescence media with 4’,6-diamidino-2-phenylindole (DAPI) (Invitrogen, P36931).

### Image acquisition and processing

Images were acquired on an inverted Zeiss 700 confocal microscope equipped with a motorized stage, 405/488/561 laser lines, and a 40X/1.3NA lens; or on an inverted Leica Thunder imaging system with individual LED lines, and 20X/0.8NA or 63 X/1.4NA lenses. Confocal images were acquired at 8-bit. All thunder images were acquired at 16-bit, subjected to computational clearing, and then set to the same fluorescence range (eg. 0-1000) within analysis cohorts before converting to 8-bit in Fiji. Representative super-resolution images in Figure 2G,2H, and 3A were acquired with 4X Super Resolution by Optical Pixel Reassignment (SoRa) on an inverted spinning disk confocal microscope (Yokogawa CSU-W1 SoRa/Nikon Ti2-E Microscope) equipped with Hamamatsu Fusion BT cameras, and 20X water (0.95 NA. WD 1.0 mm) or 60X oil (1.49 NA. WD 0.14 mm) immersion lenses. All images within analysis cohorts were acquired using identical acquisition parameters or processed identically to maintain relative differences in fluorescence, and underwent native deconvolution and denoising using NIS-Elements Software.

### Quantification of ROI fluorescence and mitochondrial size and number

The mean fluorescence, average size and number of MCU- or COX4-labeled mitochondria were quantified using the image-analysis software Fiji (v. 2.1.0/1.53c, NIH) (Schindelin et al. 2012). When feasible, all analyses were done blind to genotype or condition. A representative 100 x 100 μm ROI was selected in SO, SR and SLM of each hippocampal section analyzed. For consistency, the brightest optical Z-slice that was in focus was chosen for each ROI. Cell bodies, blood vessels and tears in the tissue were avoided as much as possible when picking an ROI. A few of the ROIs were selected at an angle (by rotating the rectangular selection tool) to avoid the tissue edge or any of the above-mentioned features.

Fiji’s *measure* tool was used to get the mean fluorescence (“mean grey value”) of the cropped ROI images. Fluorescence was measured from the cropped 8-bit images with no other adjustments made. To get the average mitochondrial size and number, individual mitochondria were segmented using “Nucleus Counter” from Fiji’s Cookbook Microscopy analysis plug-in. For the segmentation, we used an automated Otsu intensity threshold and a size threshold of 5 – 500 pixels with median 3×3 smoothing. A general “noise” threshold below which very few, if any, mitochondria were identified in the negative control images (without primary antibody) was used when necessary to adjust the automated Otsu threshold. Once the mitochondria were properly segmented, their areas were measured with the *measure* tool and exported in a .CSV file for further analysis and plotting in Python and Prism.

### Analysis of mitochondria in wild-type CA1 and CA2

40X confocal images of MCU-labeled mitochondria were segmented and quantified (as described above) from CA1 and CA2 of 3 wild-type mice, 4 hippocampal sections per mouse. 63X Thunder images of COX4-labeled mitochondria from CA1 and CA2 were similarly segmented and quantified in 7 different wild-type mice, ~4 hippocampal sections per mouse. Because MCU and COX4 were imaged at different scales on different imaging systems, mitochondrial sizes can’t be directly compared between the MCU and COX4 populations.

Background fluorescence was measured by measuring the mean fluorescence of a similar ROI from negative control images (without primary antibody) of each layer in both CA1 and CA2. The background fluorescence was subtracted from the mean fluorescence of each ROI, and all the data was normalized to the cohort average (including SO, SR and SLM of both CA1 and CA2). For plotting, the sections were averaged together to get animal averages and SR and SLM of the same animal are paired with lines.

### Analysis of Mitotag mitochondria in CA2

Five male and female Mitotag mice were perfused, horizontally sectioned, and immunostained for GFP to boost the Mitotag signal (see Immunofluorescence Methods). 16-bit images were acquired using 20X/0.8 NA and 63X/1.4 NA lenses on an inverted Leica Thunder imaging system with identical acquisition parameters per section. Images were computationally cleared and scaled to the same fluorescence intensity per section. A 100 x 100 μm ROI was selected for CA2 SO, SR and SLM regions from 63X maximum intensity projected images, then cropped, and converted to 8-bit for analysis of mitochondrial area in Fiji. Individual mitochondria were segmented using the Cookbook Microscopy plugin “Nucleus Counter” and automatically threshold using the OTSU filter. The filter set “Watershed” was applied to represent the thresholding of the original ROI most accurately. All images were processed under 5×5 median smoothing settings, and a consistent particle size range (minimum:10; maximum:10,000). Mitochondria size was measured using the measuring tool as area and exported in a .CSV file for analysis. Mitochondrial area was normalized within individual mice and the average median area of mitochondria was reported across SO, SR, and SLM regions.

### Analysis of mitochondria after MCU overexpression in CA1

MCU-labeled mitochondria were quantified in SO, SR and SLM of CA1 in 6 AAV-MCU and 5 AAV-GFP control mice. A 100 X 100 μm ROI was chosen from 63X Thunder images from each dendritic layer. MCU fluorescence, mitochondrial size and mitochondrial number were measured for each ROI as described above. The AAV-MCU data was normalized to the average AAV-GFP control for each cohort (N = 2 cohorts). A two-way RM ANOVA was performed on the normalized data, pairing dendritic layers of the same animal together, to determine the overall effect of MCU overexpression and dendritic layer.

### Protein-retention Expansion Microscopy (ProExM)

40μm horizontal mouse brain sections were expanded with 4X protein expansion microscopy (ProExM) as previously described in Campbell et al. 2021. All solutions were prepared as described by Asano et al. 2018. Sections to be expanded were incubated overnight in Acryloyl-X stock/PBS (1:100, ThermoFisher, A20770) at room temperature in the dark. Following incubation, the slices were washed twice with PBS for 15 minutes each at room temperature. The gelation solution was prepared by adding 384 uL of monomer solution, 8 uL 4-Hydroxy-TEMPO inhibitor (1:200 w/v, Sigma Aldrich, 176141), 8uL TEMED accelerator (10% v/v, Sigma Aldrich, T7024), and lastly 8uL of APS initiator (10% w/v, Sigma Aldrich, 248614) for each section. Sections were incubated in the gelation solution for 30 minutes at 4C in the dark. Gelated sections were placed on gelation chambers constructed from microscope slides with coverslips as spacers. Our gelation chambers produce gels with the thickness of a single No. 1.5 coverslip (~0.15mm thick). The chambers were filled with gelation solution and allowed to incubate at 37C for 2 hours in a humidified container. Following gelation, the gelation chamber was deconstructed and the gel was carefully removed from the chamber using a coverslip and Proteinase K-free digestion solution. Gels were then placed in digestion solution containing proteinase K (8U/mL, New England BioLabs, P8107S) and digested for 4 hours at room temperature.

For the AAV experiments, gels were placed in an alkaline buffer (100 mM Tris base, 5% (w/v) Triton X-100, 1% SDS) and digested with heat using an autoclave on the liquid cycle for 1 h at 121 C, which is a milder alternative to enzymatic digestion with Proteinase K, prior to antibody staining (Campbell et al. 2021; Asano et al. 2018). Proteinase K digested gels were immunostained prior to digestion instead. Gels were stained with DAPI (Sigma Aldrich, D9542; 1:10,000 in PBS) and MT-5 (gifted by Alex Kalyuzhny at Bio-Techne; 1:20,000 in PBS) for 30 minutes at room temperature with shaking. Kalyuzhny et al developed MT-5 as a novel mitochondrial marker specifically for use in fixed tissue (Kalyuzhny et al. 2021). MT-5 is a far-red fluorescence probe (655 nm excitation/669 nm emission) that localizes in mitochondria without cross-reacting with the cell nucleus or plasma membrane, and is highly photostable. Finally, the gels were washed twice with PBS for at least 20 minutes and either stored in PBS at 4C or fully expanded in npH20 for imaging. Images of CA2 SLM dendrites were acquired with 4X SoRa using a 20X water immersion lens (0.95 NA, WD 1.0 mm) and identical acquisition parameters on an inverted spinning disk confocal microscope (see Image acquisition and processing methods).

### Scanning Electron Microscope

Mice were anesthetized with sodium pentobarbital (euthanasia solution, 150mg/kg) and perfused with ice-cold 0.1M cacodylate buffer pH 7.4 containing 2.5% glutaraldehyde, 2% paraformaldehyde with 2mM calcium chloride for 3 minutes. The brain was removed and fixed overnight at 4C in the same fixative before vibratome sectioning (Leica VT1000S) into 350-micron thick sections in the 0.1M cacodylate buffer pH 7.4 with 2mM calcium chloride. Sections were placed back in fixative for microdissection three days later. Hemisected brain sections were placed on wax paper with a drop of fixative and a 2mm x 2mm hippocampal microdissection was obtained per brain and placed back in fixative for further processing. Tissues were postfixed with 1.5 % potassium ferrocyanide plus 2% osmium tetroxide then en bloc stained with incubations in thiocarbohydrazide solution, osmium tetroxide, uranyl acetate, and Walton’s lead aspartate. Dehydration was performed by an ethanol gradient and finished in propylene oxide. Tissues were embedded in Epon 812. The embedded tissues were sectioned to 120 nm (Leica EM UC7 ultramicrotome), mounted on a silicon wafer, and imaged in a ThermoFisher Aprea Volumescope at 2nm pixel size. Three 150 x 150 μm regions of interest were obtained per section (basal, proximal, distal CA2 dendrites). Data from one representative wild type mouse are shown. This protocol was adapted from the protocol version 2 published by NCMIR at UCSD (Deerinck et al. 2022).

### Statistical analyses

A custom Python code was written to parse the segmented data from Fiji, get the average size and number of mitochondria for each ROI, and combine the segmented data with the ROI fluorescence of each ROI. Statistical analyses were done in python (v3.7.9) or Prism (Graphpad Prism 9) with an alpha of 0.05 considered significant.

## DECLARATIONS

### Authors’ contributions

Conceptualization, S.F.; Methodology, K.E.P., D.G., M.C., L.A.C., S.F.; Formal Analysis, K.E.P., D.G., M.C., L.A.C., S.F.; Investigation, K.E.P., D.G., M.C., L.A.C., S.F.; Writing – Original Draft, K.E.P., S.F.; Writing – Review & Editing, K.E.P., M.C., D.G., S.F.; Visualization, K.E.P., M.C., D.G., L.A.C., S.F.; Supervision, S.F.; Funding Acquisition, S.F.

## Acknowledgements

We thank the members of the Farris lab for providing feedback and critically reading this manuscript, as well as the Virginia Tech animal care staff and the Cell and Molecular Imaging Core for their support.

---

**Supplemental Figure 1:**
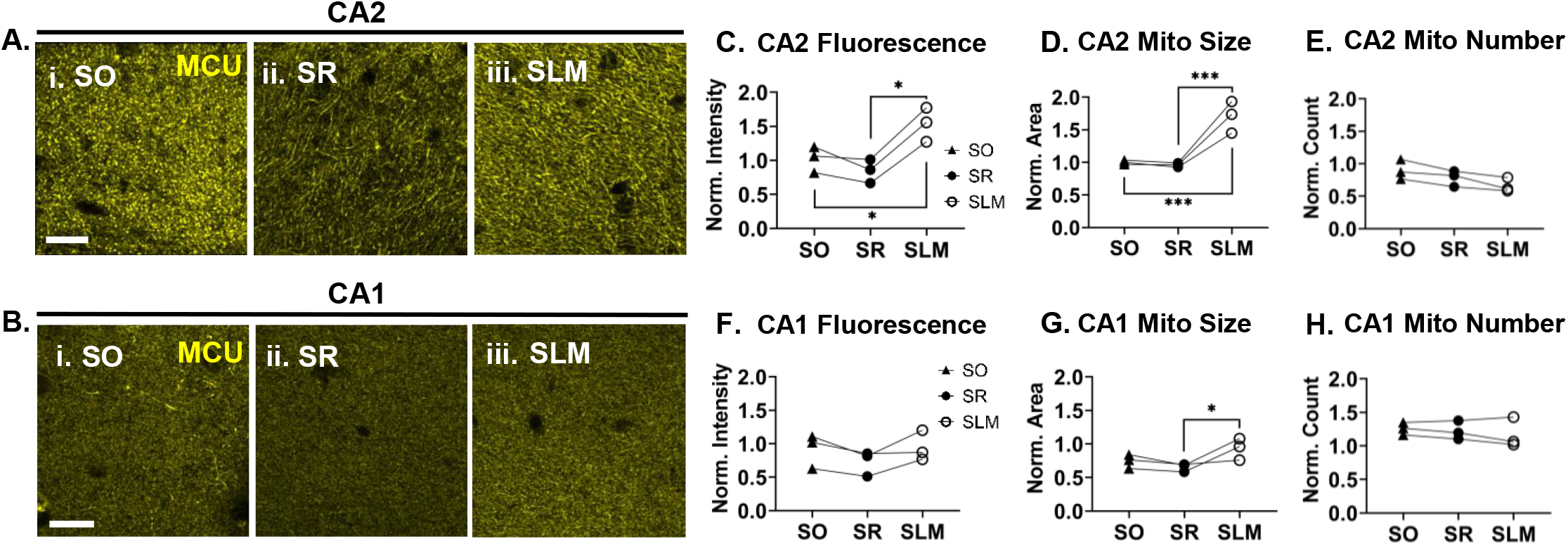
MCU staining reveals subregion and layer-specific differences in mitochondrial size and shape. **A.** Representative 40X confocal images of MCU staining in SO (*i*.), SR (*ii*.) and SLM (*iii*.) of CA2 in the same wildtype mouse as in Figure 1B-C. Images of SR and SLM are the same images used in Figure 1. **B.** Representative 40X confocal images of MCU staining in SO (*i*.), SR (*ii*.) and SLM (*iii*.) of CA1 in the same wild-type mouse. **C-E.** Quantification of MCU mean fluorescence **(C),** MCU-labeled mitochondrial size **(D),** and mitochondrial count **(E)** in all three dendritic layers of CA2. Same dataset as in Figure 1, with the addition of SO. Data is normalized to the cohort average. N = 3 animals; 4-5 hippocampal sections / animal. Stats are from the same 2way RM ANOVA as in Figure 1 and Supplemental Table 1. **F-G.** Quantification of MCU mean fluorescence **(F)**, MCU-labeled mitochondrial size **(G)**, and mitochondrial count **(H)** in all three dendritic layers of CA1 in the same hippocampal sections as **C-E**. Stats are the same as in Figure 1 and Supplemental Table 1. SO = Stratum Oriens; SR = Stratum Radiatum; SLM = Stratum Lacunosum Moleculare. Sidak’s post hoc: * = P < 0.05; ** = P < 0.01; *** = P < 0.001. Scale for A-B = 20 μm.

**Supplemental Figure 2:**
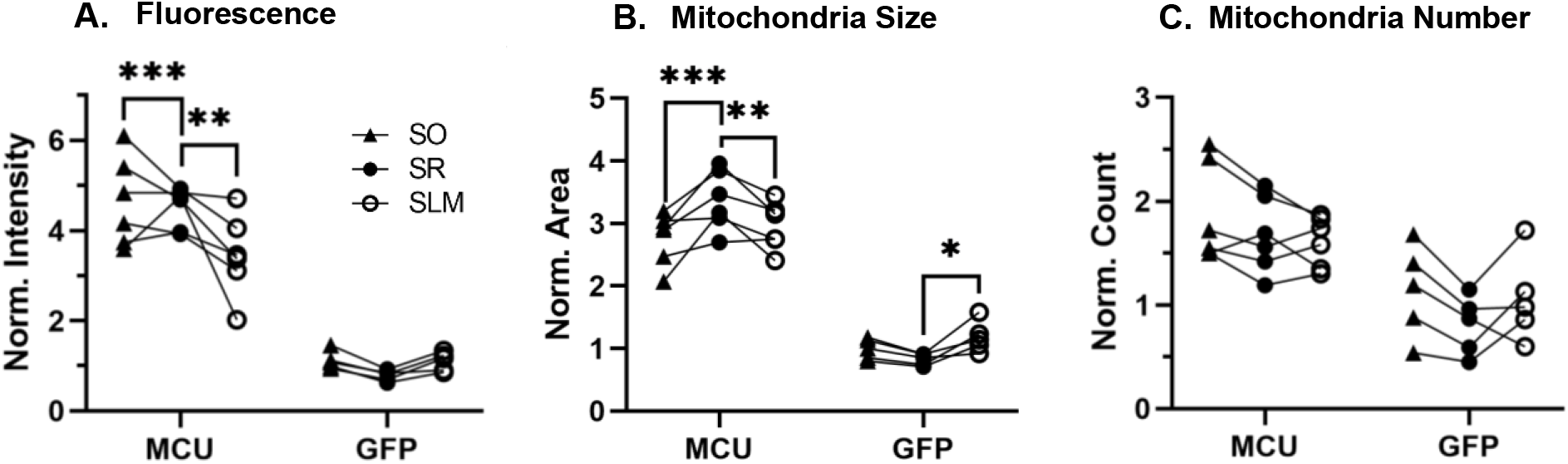
CA1 MCU overexpression increases mitochondrial size and influences layer-specific dendritic distribution. **A.** Mean MCU fluorescence of a 100 X 100 μm ROI from SO, SR and SLM of the AAV-MCU and AAV-GFP mice from Figure 4 normalized to the overall control average. Two-way RM ANOVA with Sidak’s post hoc comparing dendritic layers. N = 6 AAV-MCU mice and 5 AAV-GFP controls. Stats are the same as in Figure 4 and supplemental Table 1. **B.** MCU-labeled mitochondrial area in SO, SR and SLM of the AAV-MCU and AAV-GFP mice normalized to the overall control average. Two-way RM ANOVA with Sidak’s post hoc. **C.** Mitochondrial number in SO, SR and SLM of the AAV-mice and AAV-GFP mice normalized to the overall control average. Two-way RM ANOVA with Sidak’s post hoc. SO = Stratum Oriens; SR = Stratum Radiatum; SLM = Stratum Lacunosum Moleculare. Sidak’s post hoc: * = P < 0.05; ** = P < 0.01; *** = P < 0.001.

