## Supplemental Table 1 for "Layer-specific mitochondrial diversity across hippocampal CA2 dendrites"

| Data Set | Statistical Test | Confidence |
|---|---|---|
| **<u>MCU WT Fluorescence</u>**<br>Fig 1D<br>Fig S1 C and F | Two-way RM ANOVA<br>Sidak's multiple comparisons | <u>ANOVA</u><br>Subregion: $F_{(1,2)}$ = 309.8; P = 0.003<br>Layer: $F_{(2,4)}$ = 24.5; P = 0.006<br>Subregion X Layer: $F_{(2,4)}$ = 13.4; P = 0.017 |
| | | <u>Post Hoc comparisons</u><br>CA2 SR vs SLM: P = 0.011<br>CA2 SO vs SLM: P = 0.036<br>CA1 SR vs SLM: P = 0.473<br>CA2 SLM vs CA1 SR: P = 0.006<br>CA2 SLM vs CA1 SO: P = 0.017<br><br>CA2 SR vs CA1 SR: P = 0.951<br>CA2 SLM vs CA1 SLM: P = 0.021<br><br>*All other comparisons (15 tests) P > 0.05* |
| **<u>MCU WT Area</u>**<br>Fig 1E<br>Fig S1 D and G | Two-way RM ANOVA<br>Sidak's multiple comparisons | <u>ANOVA</u><br>Subregion: $F_{(1,2)}$ = 191.0; P = 0.005<br>Layer: $F_{(2,4)}$ = 18.0; P = 0.01<br>Subregion X Layer: $F_{(2,4)}$ = 62.1; P = 0.001 |
| | | <u>Post Hoc comparisons</u><br>CA2 SR vs SLM: P = 0.0005<br>CA2 SO vs SLM: P = 0.0006<br>CA1 SR vs SLM: P = 0.023<br>CA2 SLM vs CA1 SR: P = 0.0001<br>CA2 SLM vs CA1 SO: P = 0.0002<br>CA2 SO vs CA1 SR: P = 0.0098<br><br>CA2 SO vs CA1 SO: P = 0.032<br>CA2 SR vs CA1 SR: P = 0.016<br>CA2 SLM vs CA1 SLM: P = 0.0004<br><br>*All other comparisons (15 tests) P > 0.05* |

| | | |
|---|---|---|
| **<u>MCU WT Count</u>**<br>Fig 1F<br>Fig S1 E and H | Two-way RM ANOVA<br>Sidak's multiple comparisons | <u>ANOVA</u><br>Subregion: $F_{(1,2)}$ = 515.8; P = 0.002<br>Layer: $F_{(2,4)}$ = 12.4; P = 0.019<br>Subregion X Layer: $F_{(2,4)}$ = 2.06; P = 0.242 |
| | | <u>Post Hoc comparisons</u><br>CA2 SR vs SLM: P = 0.742<br>CA1 SR vs SLM: P = 0.999<br>CA2 SR vs CA1 SLM: P = 0.029<br>CA2 SR vs CA1 SO: P = 0.013<br>CA2 SLM vs CA1 SR: P = 0.007<br>CA2 SLM vs CA1 SO: P = 0.006<br><br>CA2 SO vs CA1 SO: P = 0.040<br>CA2 SR vs CA1 SR: P = 0.018<br>CA2 SLM vs CA1 SLM: P = 0.01<br><br>*All other comparisons (15 tests) P > 0.05* |
| **<u>COX2 WT Fluorescence</u>**<br>Fig 2D | Two-way RM ANOVA<br>Sidak's multiple comparisons | <u>ANOVA</u><br>Subregion: $F_{(1,6)}$ = 0.71; P = 0.432<br>Layer: $F_{(1,6)}$ = 13.4; P = 0.01<br>Subregion X Layer: $F_{(1,6)}$ = 16.4; P = 0.007 |
| | | <u>Post Hoc comparisons</u> (2 tests)<br>CA2 SR vs SLM: P = 0.0006<br>CA1 SR vs SLM: P = 0.0922 |
| **<u>COX2 WT Area</u>**<br>Fig 2E | Two-way RM ANOVA<br>Sidak's multiple comparisons | <u>ANOVA</u><br>Subregion: $F_{(1,6)}$ = 1.03; P = 0.349<br>Layer: $F_{(1,6)}$ = 15.5; P = 0.008<br>Subregion X Layer: $F_{(1,6)}$ = 0.12; P = 0.738 |
| | | <u>Post Hoc comparisons</u> (2 tests)<br>CA2 SR vs SLM: P = 0.002<br>CA1 SR vs SLM: P = 0.003 |

| | | |
|---|---|---|
| **<u>COX2 WT Count</u>**<br>Fig 2F | Two-way RM ANOVA<br>Sidak's multiple comparisons | <u>ANOVA</u><br>Subregion: $F_{(1,6)}$ = 0.42; P = 0.539<br>Layer: $F_{(1,6)}$ = 0.29; P = 0.612<br>Subregion X Layer: $F_{(1,6)}$ =10.4; P = 0.018<br><br><u>Post Hoc comparisons</u><br>*No post hoc comparisons made as overall effect was not significant.* |
| **<u>MitoTag Area</u>**<br>Fig 3C | One-way RM ANOVA<br>Tukey's multiple comparisons<br>(Geisser-Greenhouse correction) | <u>ANOVA</u><br>Layer: $F_{(1.6,6.5)}$ = 23.47; P = 0.001<br><br><u>Post Hoc comparisons</u> (3 tests)<br>SR vs SLM: P = 0.007<br>SO vs SLM: P = 0.009<br>SR vs SO: P = 0.234 |
| **<u>MCU OE Fluorescence</u>**<br>Fig 4D | Two-way RM ANOVA<br>Sidak's multiple comparisons | <u>ANOVA</u><br>Treatment: $F_{(1,9)}$ = 113.4; P = < 0.0001<br>Layer: $F_{(2,18)}$ = 4.8; P = 0.021<br>Treatment X Layer: $F_{(2,18)}$ = 7.24; P = 0.005<br><br><u>Post Hoc comparisons</u><br><br>**AAV-MCU vs GFP-AAV**<br>SO: P = <0.0001<br>SR: P = <0.0001<br>SLM: P = <0.0001<br><br>**Dendritic Layer**<br>AAV-MCU SR vs SLM: P = 0.002<br>AAV-MCU SO vs SLM: P = 0.0009<br>AAV-GFP SR vs SLM: P = 0.631<br><br>*All other comparisons (6 tests) P > 0.05* |

| | | |
|---|---|---|
| **MCU OE Area**<br>Fig 4C and E | Two-way RM ANOVA<br>Sidak's multiple comparisons | ANOVA<br>Treatment: $F_{(1,9)}$ = 124.4; P = < 0.0001<br>Layer: $F_{(2,18)}$ = 3.8; P = 0.041<br>Treatment X Layer: $F_{(2,18)}$ = 13.8; P = 0.0002 |
| | | Post Hoc comparisons (3 tests)<br><br>**AAV-MCU vs GFP-AAV**<br>SO: P = <0.0001<br>SR: P = <0.0001<br>SLM: P = <0.0001<br><br>**Dendritic Layer**<br>AAV-MCU SR vs SLM: P = 0.005<br>AAV-MCU SR vs SO: P = 0.0002<br>AAV-GFP SR vs SLM: P = 0.030<br><br>*All other comparisons (6 tests) P > 0.05* |
| **MCU OE Count**<br>Fig 4C and F | Two-way RM ANOVA<br>Sidak's multiple comparisons | ANOVA<br>Treatment: $F_{(1,9)}$ = 12.03; P = < 0.007<br>Layer: $F_{(2,18)}$ = 4.6; P = 0.024<br>Treatment X Layer: $F_{(2,18)}$ = 1.63; P = 0.224 |
| | | Post Hoc comparisons (3 tests)<br><br>**AAV-MCU vs GFP-AAV**<br>SO: P = 0.011<br>SR: P = 0.002<br>SLM: P = 0.07<br><br>**Dendritic Layer**<br>AAV-MCU SR vs SLM: P = 0.933<br>AAV-GFP SR vs SLM: P = 0.191<br><br>*All other comparisons (6 tests) P > 0.05* |
